# Comparing Linear and Nonlinear Finite Element Models of Vertebral Strength Across the Thoracolumbar Spine: A Benchmark from Density-Calibrated Computed Tomography

**DOI:** 10.1101/2025.04.19.649449

**Authors:** Matthias Walle, Bryn E. Matheson, Steven K. Boyd

**Affiliations:** McCaig Institute for Bone and Joint Health, University of Calgary, 3280 Hospital Drive NW, Calgary, AB, T2N 4Z6, Canada; Department of Biomedical Engineering, Schulich School of Engineering, University of Calgary, Calgary, AB T2N 1N4, Canada; Department of Radiology, University of Calgary, Calgary, AB T2N 1N4, Canada

**Author notes:** Address for correspondence: Steven K. Boyd, PhD, McCaig Institute for Bone and Joint Health, University of Calgary, 3280 Hospital Drive NW, Calgary, AB, T2N 4Z6, Canada a.

**Keywords:** Quantitative Computed Tomography, Finite Element Analysis, Vertebral Strength, Phantomless Calibration, Biomechanical Modeling, Spine, Bone Mineral Density, Opportunistic CT, Vertebral Fracture Risk, Open Benchmark Dataset

## Abstract

Opportunistic assessment of vertebral strength from clinical computed tomography (CT) scans holds substantial promise for fracture risk stratification, yet variability in calibration methods and finite element (FE) modeling approaches has led to limited comparability across studies. In this work, we provide a publicly available benchmark dataset that supports standardized biomechanical analysis of the thoracic and lumbar spine using density-calibrated CT data. We extended the VerSe 2019 dataset to include phantomless quantitative CT calibration, automated vertebral substructure segmentation, and vertebral strength estimates derived from both linear and nonlinear FE models. The cohort comprises 141 patients scanned across five CT systems, including contrast-enhanced protocols. Phantomless calibration was performed using automatically segmented tissue references and validated against synchronous calibration phantoms in 17 scans. To evaluate model performance, we implemented a nonlinear elastoplastic FE model and compared it to two linear estimates. A displacement-calibrated linear model (0.2% axial strain) demonstrated excellent agreement with nonlinear failure loads (R = 0.96; mean difference = -0.07 kN), while a stiffness-based approach showed similarly strong correlation (R = 0.92). We evaluated vertebral strength at all thoracic and lumbar levels, enabling level-wise normalization and comparison. Strength ratios revealed consistent anatomical trends and identified T12 and T9 as reliable alternatives to L1 for opportunistic screening and model standardization. All calibrated scans, segmentations, software, and modeling outputs are publicly released, providing a benchmark resource for validation and development of FE models, radiomics tools, and other quantitative imaging applications in musculoskeletal research.

## Background

Opportunistic use of clinical computed tomography (CT) offers a powerful and scalable approach to assess bone strength in routine care. These scans, often acquired for non-musculoskeletal indications, contain rich structural information that can be repurposed to assess bone health using computational methods [1, 2]. This is particularly relevant for the spine, a key skeletal site for the diagnosis and management of osteoporosis [3]. Finite element (FE) analysis applied to these images enables subject-specific simulation of mechanical loading, producing direct estimates of vertebral strength that account for both bone density and geometry [4–6]. This approach has the potential to improve risk stratification, support clinical decision-making, and expand access to bone health monitoring without requiring additional imaging or radiation exposure [7].

While several studies have advanced finite element (FE) techniques toward clinical application by linking bone strength with vertebral fracture risk, broader utility remains limited [8–11]. There remains a need for open-source and transparent modeling pipelines that can be flexibly applied across the thoracolumbar spine and tailored to diverse research questions. While tools such as biomechanical CT (O.N. Diagnostics) offer valuable FDA-approved bone strength estimates [12], improving accessibility and transparency remains essential for ensuring broader population representation. In parallel, differences in FE implementation introduce further variability. Nonlinear models, which incorporate assumptions about yield properties, asymmetry, and post-yield behavior, aim to reflect bone failure mechanics more realistically—but rely on parameters that are difficult to validate and may change with aging or disease. Linear models are more reproducible and efficient but may oversimplify failure behavior [4]. Despite these trade-offs, few studies have directly compared linear and nonlinear FE analysis across vertebral levels. Furthermore, while clinical imaging often captures multiple vertebrae, research remains heavily focused on the first lumbar vertebra (L1), restricting insight into regional variation in strength and complicating the use of consistent thresholds across the spine.

To address these limitations, this study systematically compares vertebral strength estimates from linear and nonlinear FE analyses across the thoracic and lumbar spine. We further quantify intervertebral strength ratios to evaluate whether previously established strength thresholds can be extrapolated beyond L1, with the aim of supporting standardized multilevel assessments. All density-calibrated scans, vertebral segmentations, and strength estimates are made publicly available to facilitate reproducibility and future method development. These resources cover all thoracic and lumbar vertebrae (T1–L5) and are intended to fill a critical gap in vertebral strength modeling, enabling transparent comparison of FE analysis pipelines across research groups.

### Data Description

This study provides a benchmark resource for vertebral strength estimation from density-calibrated clinical CT scans. It builds upon the publicly available VerSe 2019 dataset [13–15] by incorporating phantomless calibration for extracting quantitative CT measurements, standardized vertebral substructure segmentations, and strength estimates from both linear and nonlinear FE models (**Fig. 1**). Nonlinear FE simulations are computationally expensive and often require substantial computational infrastructure, making them impractical for routine use. By providing these outputs, we offer a ready-to-use reference for future studies aiming to develop and validate simplified or surrogate models—without the burden of reprocessing high-fidelity simulations. The dataset includes individually calibrated and aligned vertebrae with accompanying segmentation masks, ensuring that all inputs are standardized and spatially harmonized. This allows researchers to convert the provided data directly into custom finite element models without introducing variability from preprocessing steps such as segmentation, orientation, cropping, or resampling, which could otherwise confound comparative analyses [16]. Further, all software for density calibration and finite element analysis has been made publicly available under open-source licenses to allow researchers to build on these results in their own datasets. The goal of this extended dataset is to support reproducible and standardized biomechanical analysis of the thoracic and lumbar spine. It enables consistent comparison of linear and nonlinear FE modeling approaches, supports the development of machine learning and radiomics tools, and facilitates investigations into vertebral strength variation across spinal levels. In addition, the dataset can be used to benchmark calibration techniques and to support opportunistic CT-based assessment of bone health in clinically acquired scans.

**Figure 1.**
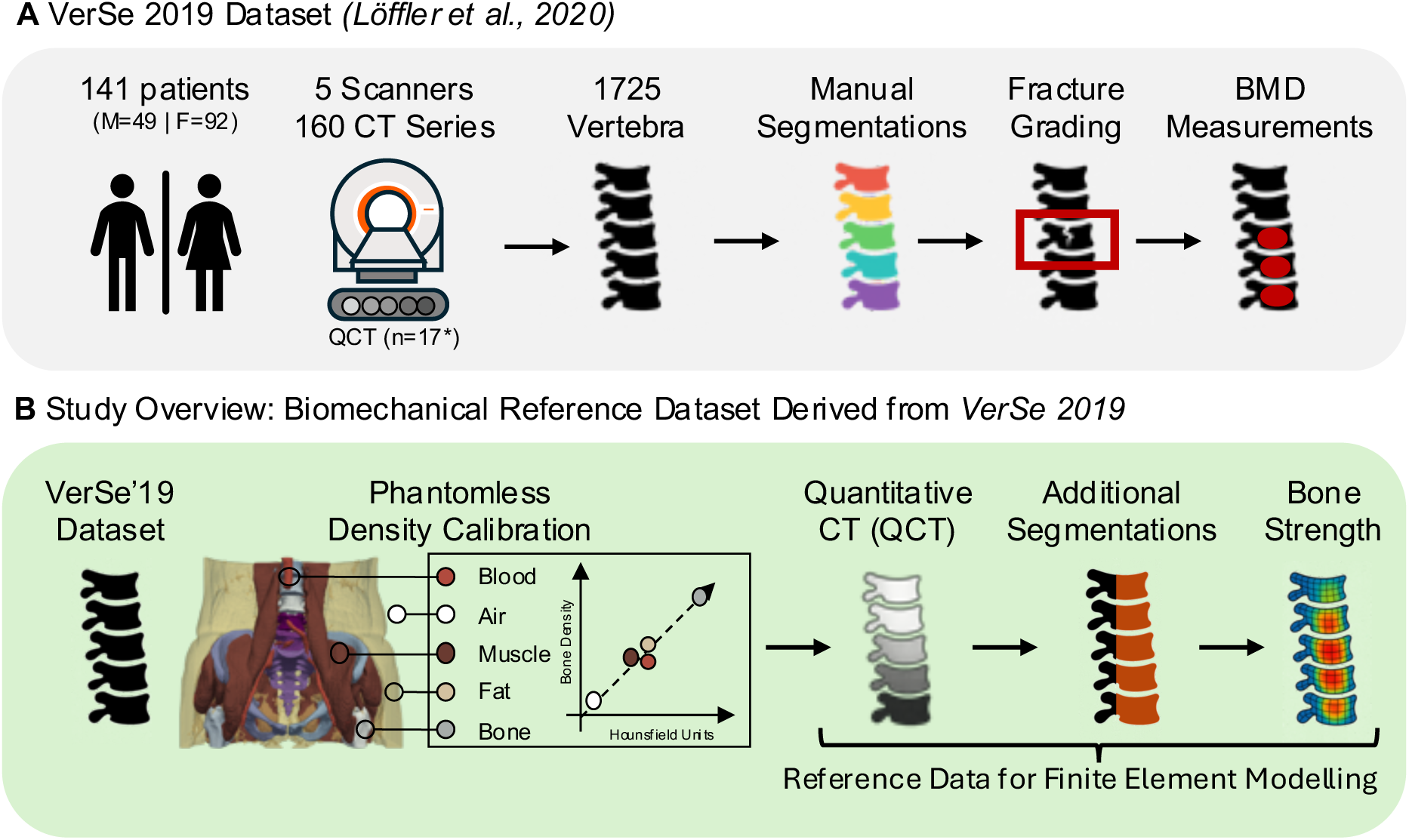
Overview of the VeSe 2019 dataset and its extension in this study. **[A]** The original VerSe 2019 dataset includes CT scans of 141 patients (49 male, 92 female) acquired across five scanners, resulting in 1725 annotated vertebrae. Annotations include vertebral level labels, manual segmentations, Genant-based fracture grading, and volumetric bone mineral density (vBMD) measurements. **[B]** In this study, we derived a biomechanical reference dataset from VeSe 2019 by applying phantomless density calibration to convert CT scans into density-calibrated quantitative CT (QCT) scans. These scans were used to generate additional segmentations required for finite element modeling, including segmentations of the vertebral body and spinal processes. Bone strength was estimated using linear and nonlinear finite element modeling at all vertebral levels. All data and derived outputs will be made publicly available to support benchmarking and reproducibility in computational spine research.

### Analyses

#### Phantomless calibration enables consistent estimation of Bone Mineral Density

To enable density-based finite element modeling, we calibrated CT-derived Hounsfield Units (HU) to volumetric bone mineral density (vBMD) using a phantomless approach [17]. A total of 17 scans in the VerSe dataset (**Table 1**) included a physical calibration phantom, allowing comparison of phantomless calibration to both synchronous and asynchronous phantom calibration methods (**Fig 2**). Phantomless calibration showed strong agreement with synchronous phantom calibration (R = 0.91; **Fig. 3A**). Bland–Altman analysis revealed a small mean difference of 1.2 mg/cm³ and narrow limits of agreement (**Fig. 3B**). Agreement was consistent across contrast-enhanced scans, including both portal venous (ce-pv) and arterial (ce-art) phases. Comparison with scan-wise asynchronous calibration values from Loffler, Sekuboyina [13] resulted in lower agreement (R = 0.72; **Fig. 3C**) and a larger mean offset of –29.5 mg/cm³ (**Fig. 3D**).

**Figure 2.**
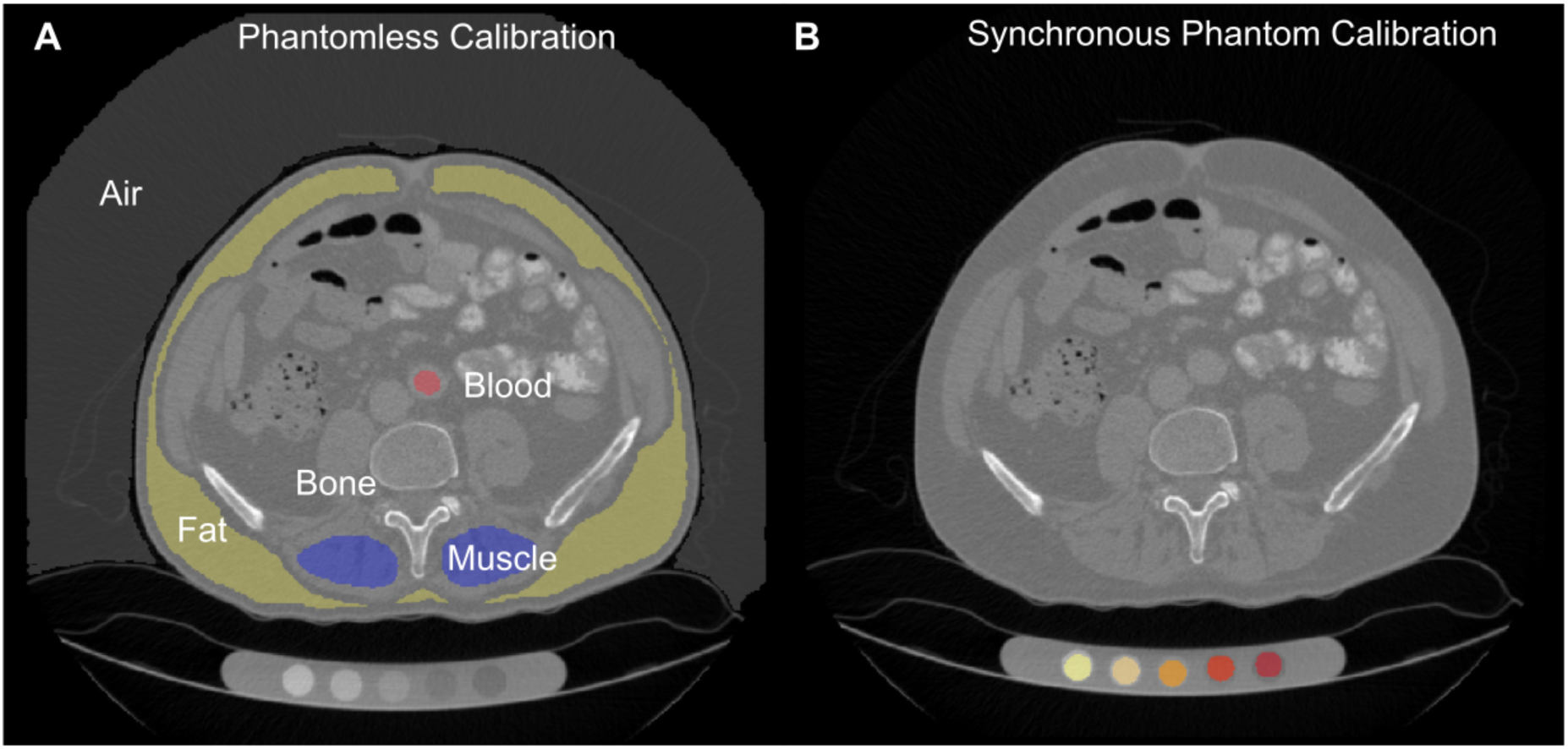
Phantomless versus synchronous phantom calibration. **[A]** Example of phantomless calibration based on tissue-equivalent regions of interest including air, fat, muscle, blood, and bone. **[B]** Example of synchronous phantom calibration using a physical calibration phantom scanned with the patient.

**Figure 3.**
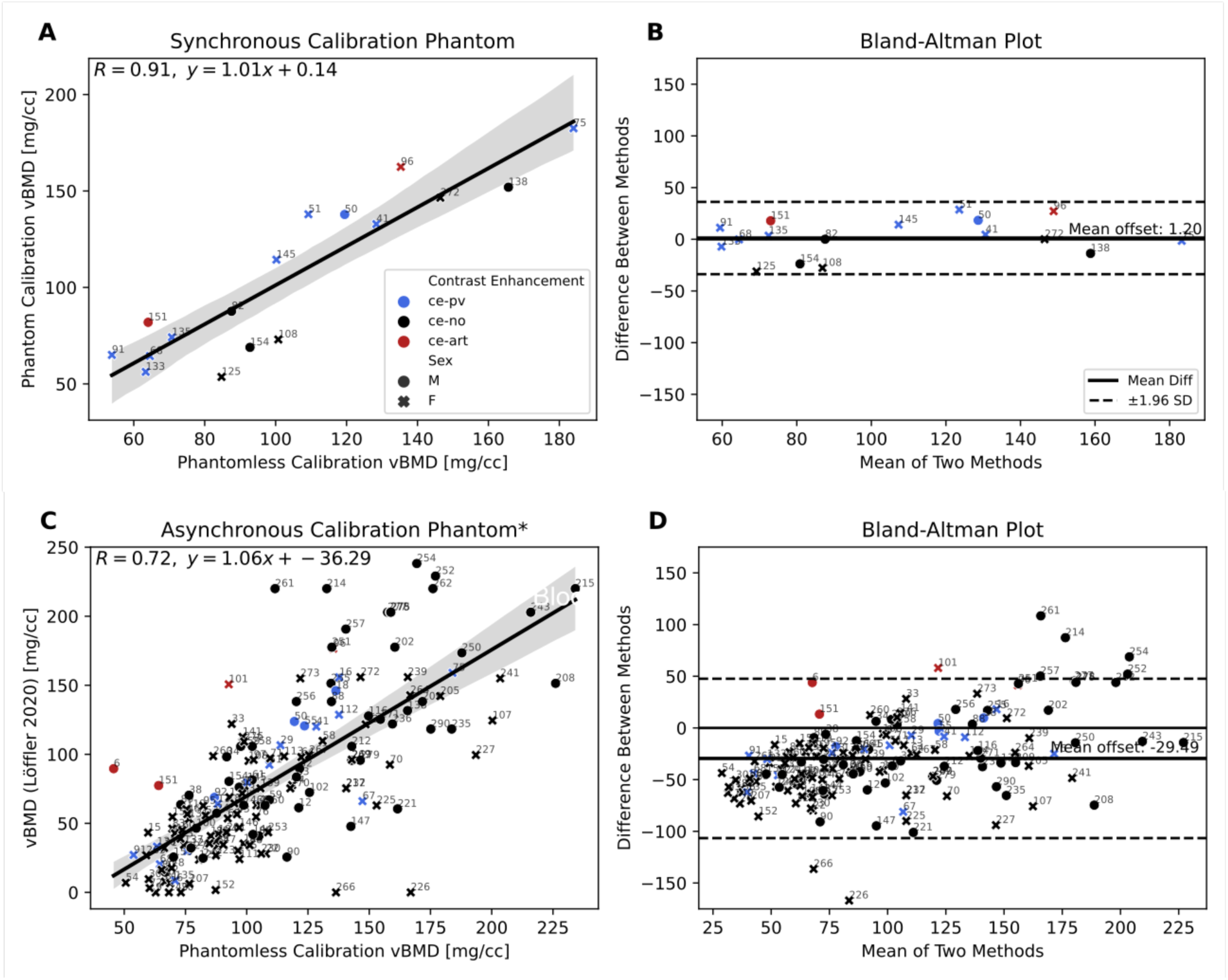
Comparison of vBMD measurements across calibration methods. **[A]** Correlation between phantomless calibration and synchronous phantom calibration (n=17) shows strong agreement (R = 0.91). VerSe scan IDs are labeled on each image to indicate scan-wise correspondence. **[B]** Bland–Altman plot indicating small mean bias and narrow limits of agreement between methods. **[C]** Comparison between phantomless and asynchronous calibration (n=141, from Löffler et al., 2020) shows lower agreement (R = 0.72). *Lower agreement is likely due to unreported exact measurement region in Löffler et al., 2020. **[D]** Bland–Altman plot for asynchronous calibration shows larger mean offset and wider variation.

**Table 1.**
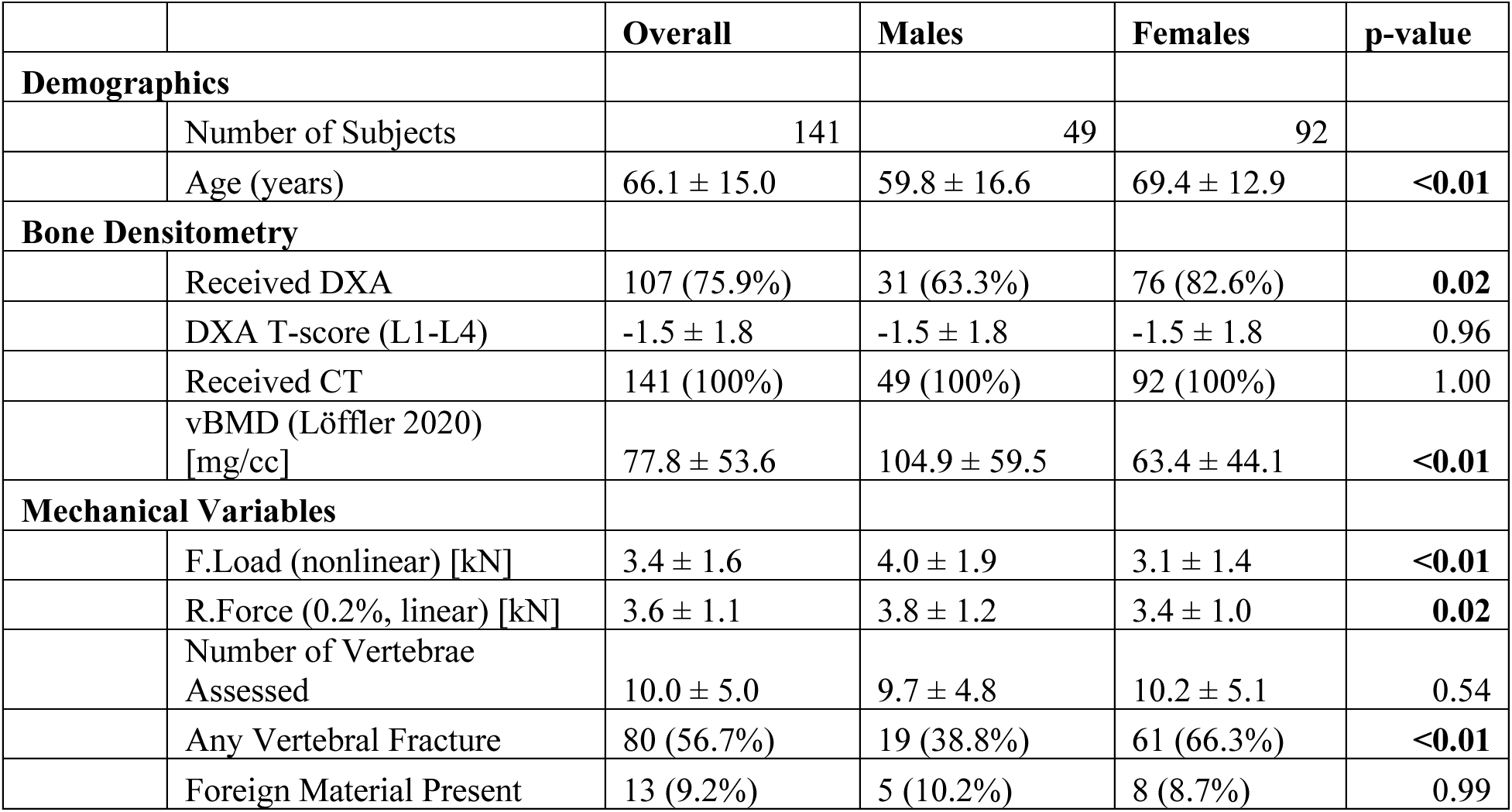
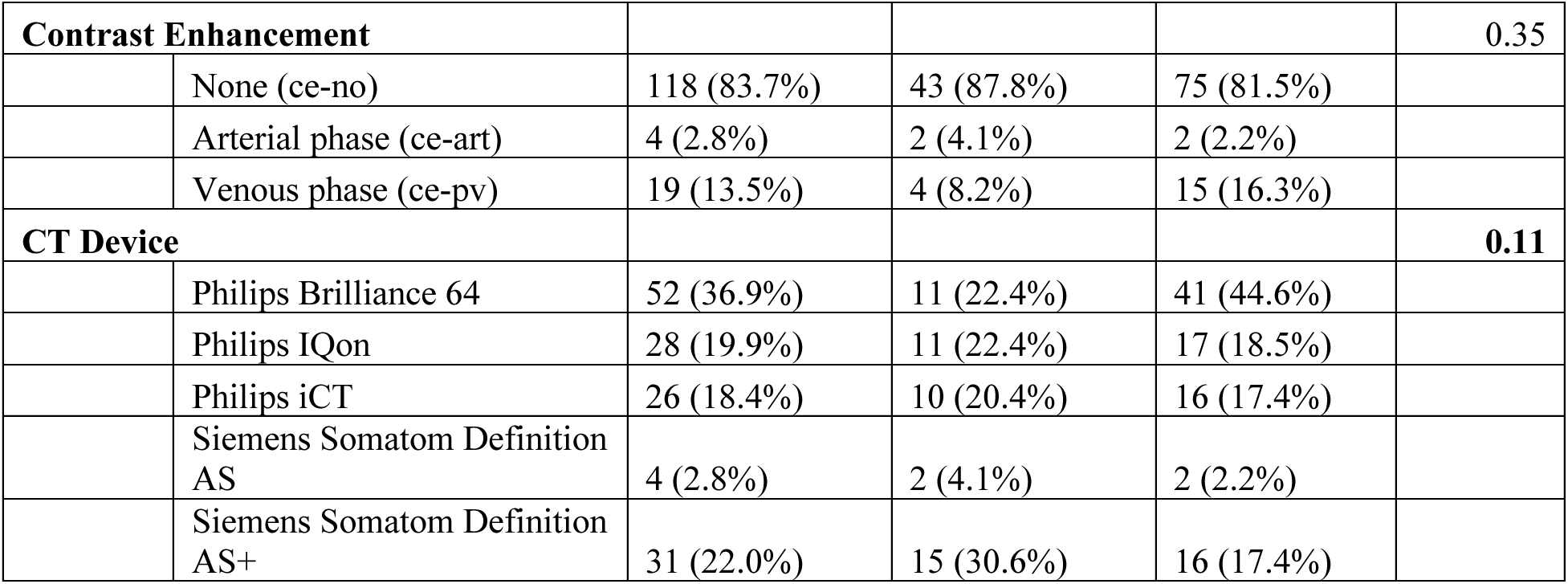
Participant characteristics and imaging-derived measures including dual X-ray absorptiometry (DXA), volumetric bone mineral density (vBMD), and finite element analysis derived failure load (F.Load) and reaction forces (R.Force), stratified by sex. Values are presented as mean ± standard deviation or count (percentage). Group differences between males and females were assessed using t-tests or chi-squared tests, as appropriate.

#### Linear models approximate nonlinear vertebral strength estimates

We evaluated two linear finite element (FE) modeling approaches to approximate vertebral failure load and compared their predictions against nonlinear FE simulations.

In the first approach, we developed a displacement-calibrated method using nonlinear simulation data. Nonlinear force–displacement curves were generated for three representative samples with low, medium, and high vBMD (**Fig. 4**). Across these cases, a displacement threshold of 0.2% was found to produce linear reaction forces (R.Force) closely matching nonlinear failure loads. Applying this calibrated threshold to the full cohort resulted in a strong correlation between linear and nonlinear strength estimates (R = 0.96, **Fig. 5A**), with a mean difference of –0.07 kN (**Fig. 5B**). Importantly, the few outliers where linear analysis overestimated strength occurred only in individuals with high strength who are not considered at risk.

**Figure 4.**
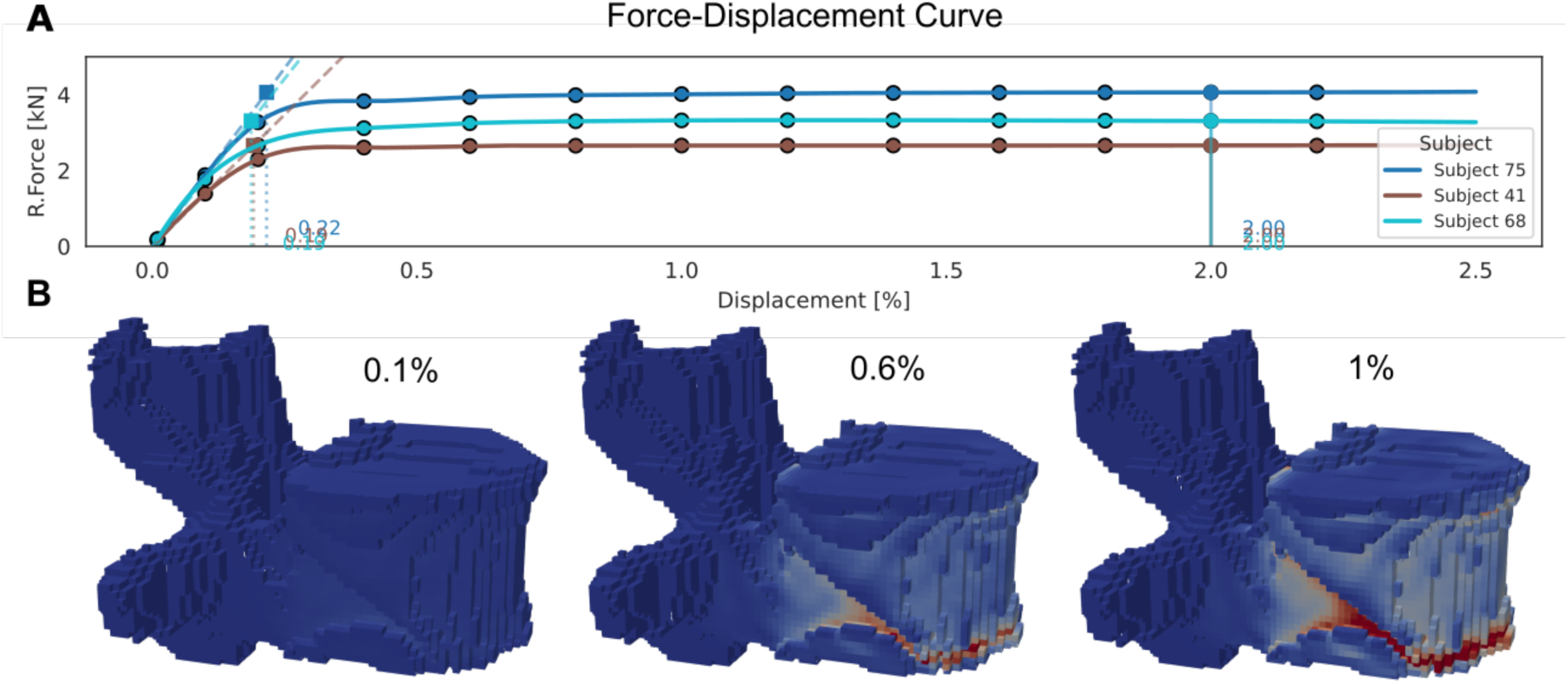
Finite element force estimation with phantomless calibration. **[A]** Reaction force–displacement curves from nonlinear simulations in three representative vertebrae (VerSe subjects 41, 68, and 75). Dashed vertical lines indicate where a linear fit to the initial slope reaches the failure load, defined as the total reaction force at 2% displacement (solid vertical line). **[B]** Corresponding Strain distributions at increasing displacement levels (0.1%, 0.6%, 1.0%) for subject 41 at the L1 vertebra. Shades of red indicate higher strain; shades of blue indicate lower strain.

**Figure 5.**
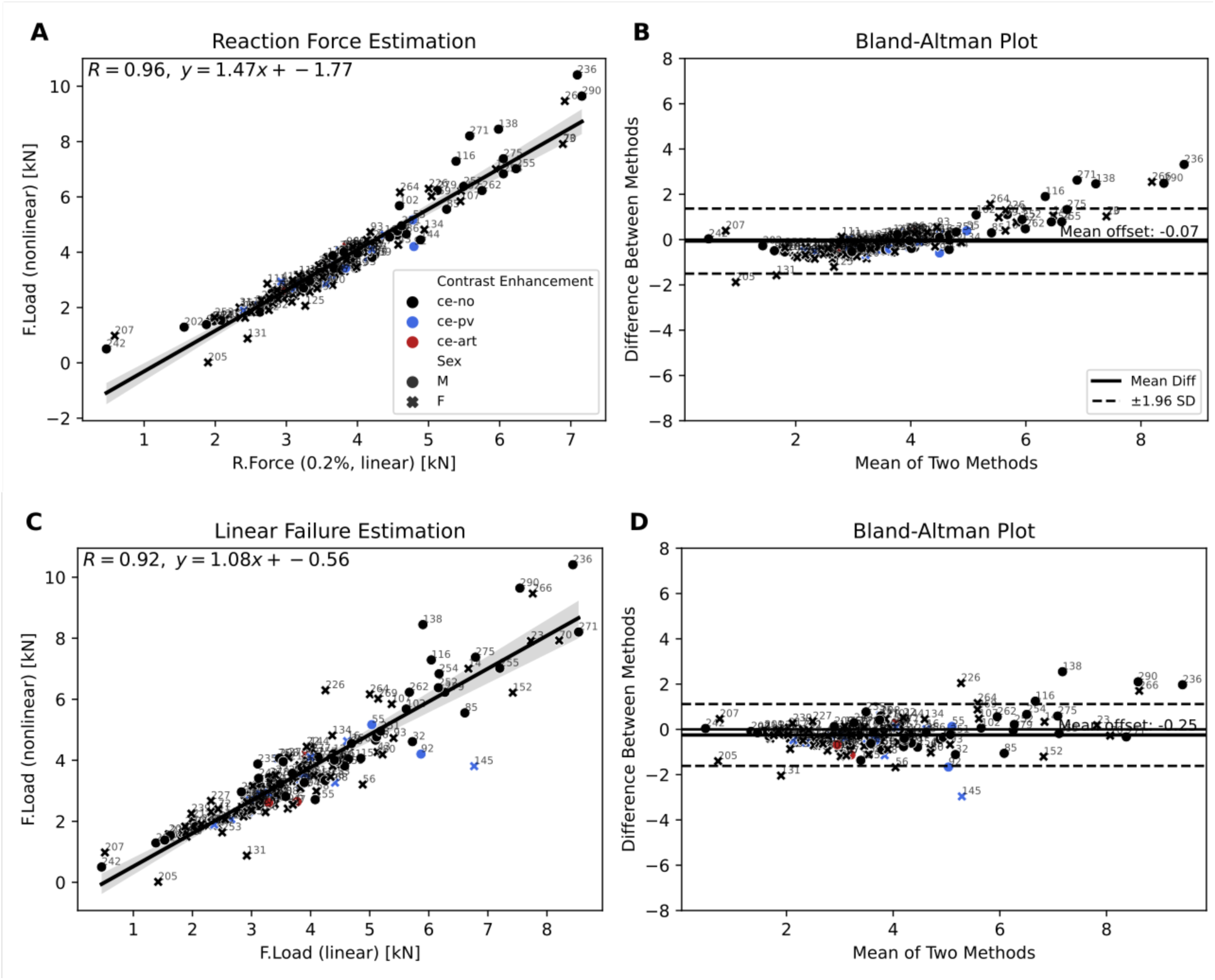
Linear vs nonlinear failure estimation. **[A]** Correlation between linear and nonlinear failure load estimates shows strong agreement (R = 0.96). VerSe scan IDs are labeled on each image to indicate scan-wise correspondence. **[B]** Bland–Altman plot confirms high agreement and minimal bias between the two methods. **[C]** Correlation of linear reaction force and nonlinear failure load estimates (R = 0.92). **[D]** Bland–Altman plot shows small positive bias in nonlinear load predictions.

We compared this to an approach adapted from prior work that estimates vertebral strength using a column-based linear model [18]. Specifically, compressive failure load (F.Load) was calculated as the product of model stiffness (K_FE_ and vertebral height (H), scaled by a constant yield strain factor (0.0068), under the assumption of uniform axial loading and average material failure properties across samples. This method yielded a slightly lower correlation with nonlinear failure loads (R = 0.92; **Fig. 5C**), the regression slope was closer to 1, indicating better agreement in scale. The mean difference between both methods was -0.25 kN (**Fig. 5D**).

To assess the impact of vBMD calibration on these predictions, we compared strength estimates obtained from phantomless- and phantom-based QCT images using the displacement-calibrated method at the 0.2% strain threshold. The correlation between phantom-based and phantomless-derived FE estimates was high (R = 0.70), with a mean difference of –0.01 kN, indicating that phantomless calibration does not introduce substantial bias in FE-based strength estimation.

#### Vertebral strength varies consistently across spinal levels

To enable comparison of FE-derived strength estimates across different spinal levels, we quantified vertebral strength relative to L1 using two normalization approaches. The first assessed within-subject differences between adjacent vertebrae (**Fig. 6A**), while the second used the population-average L1 strength as a reference (**Fig. 6B**).

**Figure 6.**
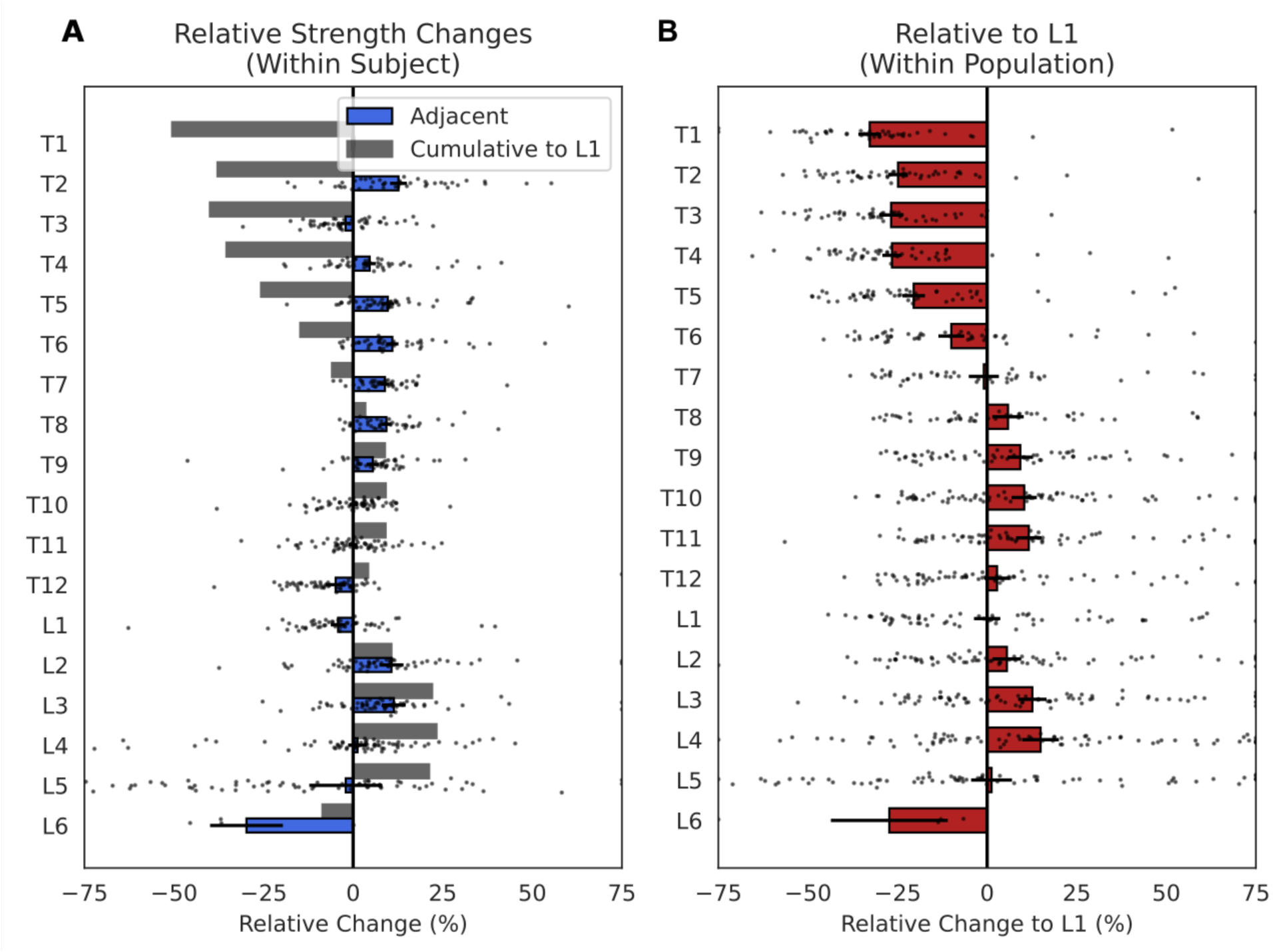
Reference maps of vertebral strength variation. **[A]** Within-subject relative strength changes show localized differences between vertebrae. **[B]** Within-population analysis relative to L1 reveals consistent patterns of regional variation across subjects, highlighting decreased strength in lower lumbar vertebrae

Both approaches revealed consistent anatomical trends, with strength decreasing towards the upper thoracic and increasing towards the lower lumbar levels. Among thoracic vertebrae, T12 (+2.2%) and T9 (-2.1%) showed the smallest deviation from L1, suggesting they are good alternative targets for opportunistic bone strength assessments. In contrast, lower lumbar vertebrae such as L2 and L3 showed strength increases of 8.5% and 13.7% relative to L1. Sex-specific analysis showed similar trends with minor variation in magnitude. For example, strength at T3 was 58.2% lower than L1 in males and 44.9% lower in females (**Table 2**).

**Table 2:**
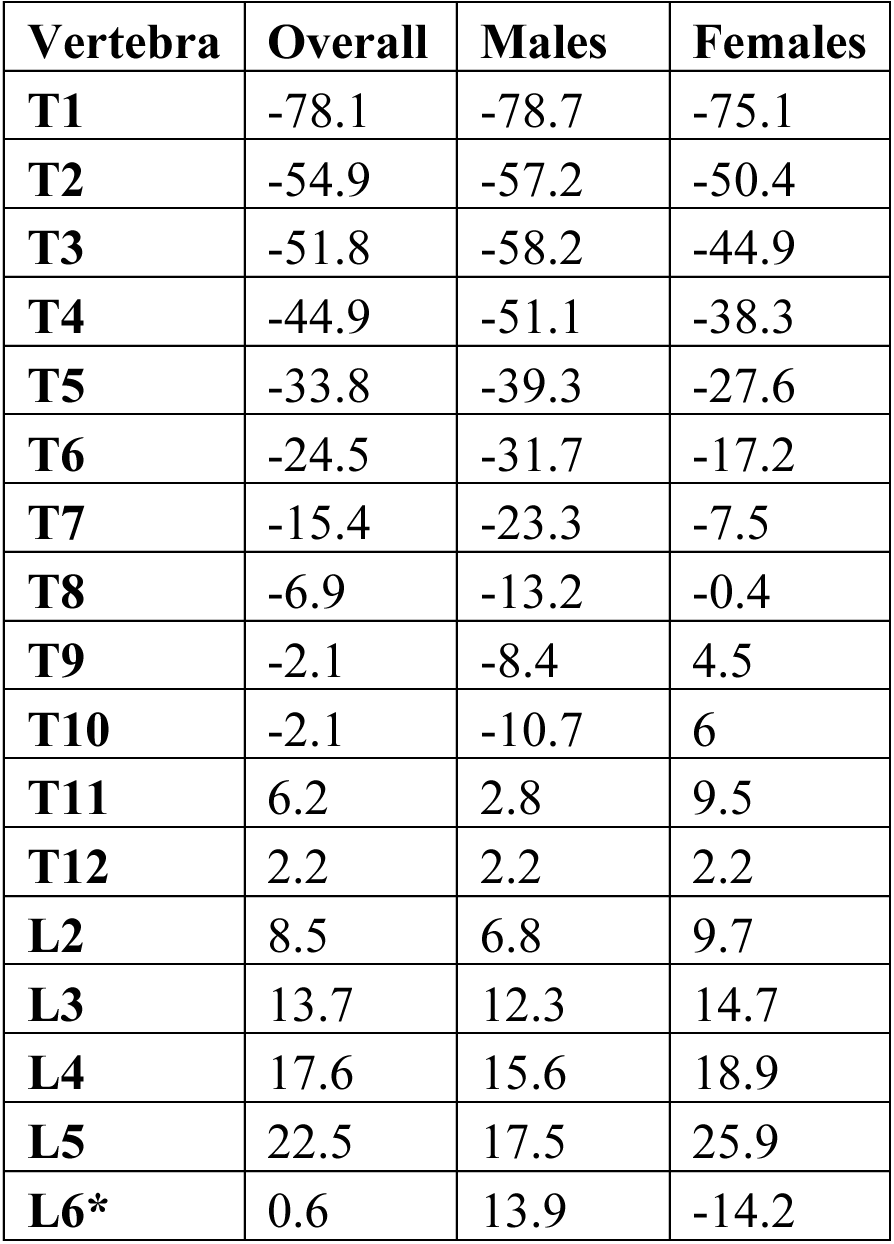
Relative vertebral strength compared to L1 across thoracic and lumbar levels. Values represent mean percentage differences in FE-derived strength estimates relative to L1, calculated separately for males and females. Negative values indicate lower strength compared to L1. *Small sample size (n=6).

| Vertebra | Overall | Males | Females |
| --- | --- | --- | --- |
| T1 | -78.1 | -78.7 | -75.1 |
| T2 | -54.9 | -57.2 | -50.4 |
| T3 | -51.8 | -58.2 | -44.9 |
| T4 | -44.9 | -51.1 | -38.3 |
| T5 | -33.8 | -39.3 | -27.6 |
| T6 | -24.5 | -31.7 | -17.2 |
| T7 | -15.4 | -23.3 | -7.5 |
| T8 | -6.9 | -13.2 | -0.4 |
| T9 | -2.1 | -8.4 | 4.5 |
| T10 | -2.1 | -10.7 | 6 |
| T11 | 6.2 | 2.8 | 9.5 |
| T12 | 2.2 | 2.2 | 2.2 |
| L2 | 8.5 | 6.8 | 9.7 |
| L3 | 13.7 | 12.3 | 14.7 |
| L4 | 17.6 | 15.6 | 18.9 |
| L5 | 22.5 | 17.5 | 25.9 |
| L6* | 0.6 | 13.9 | -14.2 |

## Discussion

This study provides a benchmark dataset for vertebral strength estimation from density-calibrated clinical CT scans, enabling direct comparison of linear and nonlinear finite element (FE) modeling approaches across the thoracic and lumbar spine. We extend the publicly available VerSe dataset with phantomless calibration, vertebral and posterior element segmentations, and strength estimates from two common QCT-based material models—creating a comprehensive, open resource for comparing and developing FE methods without bias resulting from preprocessing steps. Our findings demonstrate that a displacement-calibrated linear model can closely approximate nonlinear strength predictions and that vertebral strength varies systematically by level and sex. Importantly, we identify alternative vertebral targets such as T12 and T9 that may serve as more appropriate fallback sites when L1 is unavailable. This work supports greater standardization and reproducibility in FE-based bone strength assessment and facilitates broader applications of CT-derived biomechanics in clinical and research settings.

Our results demonstrate that a previously developed phantomless density calibration method [17, 19] can be successfully applied to an external dataset acquired across different scanners and manufacturers, showing strong agreement with synchronous phantom-based calibration. Notably, this dataset expands beyond prior work [17, 19] by including not only abdominal scans but also thoracic and lumbar spine levels, further demonstrating the generalizability of the approach. While comparisons with previously published patient-level vBMD values showed slightly lower correlations [13], these differences are likely due to variations in measurement regions and segmentation approaches. For example, while the VerSe study assessed L1–L4 using expert-drawn regions only when intact, it remains unclear how cases with fractures, missing levels, or anatomical anomalies were exactly addressed [13]. Although scanner-specific calibration equations were previously published for a subset of patients (n=15, [20]), we chose internal calibration as the reference to better account for potential scanner drifts. To support transparency and reuse, our study not only provides vBMD values but also a method to reconstruct density-calibrated images, vertebral masks, and metadata on analyzed vertebrae. This enables further downstream applications, from finite element modeling to radiomics or soft tissue analysis, and expands the utility of phantomless calibration in opportunistic CT imaging.

The estimated vertebral failure forces from our models are consistent with prior studies using similar QCT-based finite element approaches and validation cohorts [18, 21]. Although these values fall below strength thresholds reported in larger clinical studies [8], this is expected given our cohort, which includes a high prevalence of fractures (91 out of 141 subjects have at least one spinal fracture) and a large proportion of females [13]. To estimate failure forces, we implemented two widely used approaches: a linear elastic model based on the density–modulus relationship from Crawford, Cann [18], and a nonlinear model using the density-dependent yield stress formulation from Kopperdahl, Morgan [22]. Both models assign material properties voxel-wise based on apparent density from calibrated CT scans, implicitly capturing cortical bone through higher density values—given QCT’s limited resolution and partial volume effects that often overestimate cortical thickness [23]. We also provide cortical bone segmentations based on a previous approach [24] for studies interested in explicitly analyzing this compartment. While some models apply different material behavior in tension and compression [25–28], we applied a single yield stress for both. Although the spine experiences various loading modes, including shear and torsion [29], most FE studies focus on compressive loading, which primarily induces compressive stresses within the vertebral body, with minimal tensile components. Together, these modeling choices balance physiological relevance with computational efficiency and comparability. At the same time, our framework remains flexible to accommodate more detailed constitutive models for future studies, including distinct yield stress formulations for compression and tension, which may be required for other skeletal sites such as the hip.

Our findings highlight that the choice of vertebral level can significantly affect finite element-based strength estimates and should be considered in study design and interpretation. While L1 is the most commonly analyzed vertebra, L2 is often used as a fallback when L1 is unavailable. However, we found that L2 is, on average, almost 10% stronger than L1, which could lead to overestimation of strength if substituted directly. In contrast, T12 showed only a 2% difference from L1 and demonstrated consistent anatomical similarity across individuals, suggesting it may be a more appropriate alternative. Although current FDA-approved protocols for the assessment of osteoporosis frequently allow analysis of any vertebra from T12 to L3 [12], our results suggest that inter-level strength differences within this range are non-negligible and should be considered in clinical interpretation and cross-study comparisons. Chest CT scans performed for indications such as lung cancer screening or pulmonary embolism often exclude the lower thoracic and lumbar spine. Our results show that T9, which is typically included in these protocols, serves as a suitable fallback for strength estimation, with values within 2% of L1. Notably, a prior study suggested that T8 predicts fractures as well as L1, and developed vertebral strength scale factors in a small cohort (n=22), though these were not published for all vertebral levels [10]. These findings support the need for more consistent vertebral targeting and the development of level-adjusted reference values for spinal strength estimation using finite element models.

Several limitations should be considered when interpreting the results of this study. First, the VerSe dataset includes a high prevalence of vertebral fractures, reflecting a population with greater skeletal fragility. Consequently, the strength estimates derived from our finite element models may be lower than those reported in previous studies involving healthier or younger cohorts. Nonetheless, this cohort is representative of the clinical population most likely to benefit from opportunistic CT-based screening for skeletal fragility. Second, while our phantomless calibration approach showed strong agreement with in-scan density phantoms in a subset of 17 scans, the absence of full calibration certificates (data was not available on request) for all scanners may have contributed to slightly lower correlations when compared to phantom-based calibration. Further, some scans included partial phantoms within the field of view, which may have introduced additional variability. However, these effects are expected to be marginal, and our results remain consistent with previously published asynchronous calibration data [17]. Third, our finite element models include common assumptions for clinical QCT applications: voxel-wise density-based material properties without explicit cortical-trabecular separation, and standardized uniaxial boundary conditions. While these simplifications may limit anatomical and loading specificity, they enhance reproducibility across vertebral levels and modeling approaches. To support future studies aiming to investigate more complex models—including anisotropic material behavior or subject-specific boundary conditions—we provide all necessary data, segmentations, and open-source software. Lastly, although clinical fracture annotations are included, the lack of prospective follow-up limits assessment of predictive performance for future fractures.

A central strength of this study is the development and validation of a fully automated pipeline for estimating vertebral strength directly from routine clinical CT scans, without requiring in-scan calibration phantoms. This is particularly relevant for opportunistic imaging, where phantoms are rarely present and manual processing is impractical at scale. Moreover, in-scan phantoms can introduce beam hardening artifacts or degrade image quality, potentially affecting both visual interpretation and downstream analysis. By integrating phantomless calibration, vertebral segmentation, and standardized finite element modeling into a cohesive pipeline, we enable reproducible strength estimation across the thoracolumbar spine. This end-to-end approach reduces preprocessing variability and facilitates biomechanical assessment in real-world clinical datasets, supporting broader implementation and translational research.

### Potential Implications

This dataset provides density-calibrated CT images with broad applicability beyond vertebral strength and fracture risk estimation [6]. Unlike conventional CT, our calibration technique enables standardized, reproducible tissue density measurements, opening avenues for quantitative assessment of muscle quality and geometry [30], body composition including visceral and subcutaneous fat [31], and the density of calcified plaques within blood vessels that may be useful in detecting high-risk patients with coronary atherosclerosis [32]. Quantitative muscle imaging, for example, is increasingly used to detect early signs of atrophy or infiltration associated with cancer, diabetes, and renal failure [33, 34] and reflects the growing interest in the bone–muscle unit [35, 36]. Together, these capabilities support integrated musculoskeletal and cardiometabolic phenotyping, improve precision in evaluating age- and disease-related changes, and enable advanced applications such as radiomics-driven for opportunistic osteopenia and osteoporosis screening [37], or finite element modeling that rely on consistent quantitative input [38].

Crucially, while opportunistic CT has emerged as a promising avenue for large-scale health screening, there remains a lack of openly available datasets that provide internal calibration alongside phantom-calibrated CT outputs. Our resource fills this gap by offering individually calibrated scans, enabling the evaluation of internal calibration methods across a diverse range of spinal levels and patient anatomies. Further, FE modeling studies have historically been limited by the inaccessibility of original models, making it difficult to directly compare modeling pipelines or validate published thresholds for fracture risk. Our dataset addresses this challenge by providing strength estimates derived from both linear and nonlinear FE analysis, enabling transparent comparisons without requiring reimplementation of complex FE setups. This is particularly valuable for groups aiming to develop simplified surrogate models or explore new thresholds based on intervertebral strength ratios. The availability of calibrated, aligned scans with corresponding anatomical segmentations ensures that users can readily generate their own custom FE models without concerns about preprocessing variation, a common source of discrepancy in biomechanical simulations [16]. Together, these features lay the groundwork for more reproducible, scalable, and comparable computational studies for osteoporosis diagnosis and management.

## Methods

### Study Cohort and Imaging Data

This study builds upon the publicly available VerSe 2019 dataset [13–15], which includes CT scans of 141 patients (49 males, 92 females), encompassing 1725 vertebrae labeled across the thoracic and lumbar spine (T1–L6). CT scans were acquired across five different scanners (Philips and Siemens models) using standardized protocols. All scans met minimum imaging requirements of 120-kVp acquisition with sagittal reformations reconstructed using filtered back projection with a bone kernel optimized for edge detail. A spatial resolution of at least 1 mm in the craniocaudal direction was ensured to preserve anatomical fidelity for downstream segmentation and modeling. All images were acquired for clinical purposes unrelated to musculoskeletal health, resulting in a dataset enriched with incidental vertebral fractures. The scans were collected for various indications including cancer staging, exclusion of abdominal pathology, postoperative evaluation, and back pain assessment. Both non-enhanced and contrast-enhanced scans are included, with contrast administered in either the arterial or portal venous phase. Ethical approval for publication of the anonymized dataset was obtained from the Technical University of Munich (Proposal 27/19 S-SR), and the dataset was released under the Creative Commons Attribution-ShareAlike 2.0 license (CC BY-SA 2.0).

Annotations provided in the VerSe dataset include vertebral level labels, fracture grades based on Genant criteria, and manual vertebral segmentations. Segmentations were generated using a deep learning framework (Btrfly Net and U-Net architectures), followed by manual refinement by trained annotators and neuroradiologists. Final segmentations are provided in NIfTI format labeled by vertebral level, with non-bone structures such as implants and cement removed.

### Vertebral Segmentation

The original VerSe dataset included manual segmentations of whole vertebrae for each annotated level. For finite element modeling, we generated additional segmentations of the vertebral body and posterior elements. This was achieved using an in-house trained nnU-Net model [39], which was applied to the original segmentation mask to relabel substructures. The final masks were saved in NIfTI format and used in subsequent modeling and calibration steps.

### Phantomless Density Calibration

To convert CT-derived Hounsfield Units (HU) to vBMD, we applied a phantomless calibration method adapted from Michalski, Besler [17]. Instead of manually placing tissue-equivalent regions of interest, we used an in-house trained nnU-Net, trained on the publicly available TotalSegmentator dataset [40], to automatically segment key reference tissues, including subcutaneous adipose tissue, autochthonous and gluteal muscles, the aorta, the common iliac arteries, and air. Air segmentation was performed by thresholding voxels with HU values between –950 and –1050, followed by largest connected component filtering to isolate background air outside the body. Tissues were eroded by five voxels to avoid partial volume and boundary effects. HU means were extracted from these segmented regions and used to construct a scan-specific calibration curve mapping HU to equivalent K₂HPO₄ density. For vBMD measurement, we applied a five-voxel erosion to the vertebral body mask to reduce partial volume effects and minimize the influence of the dense cortical shell. In total, 17 scans in the dataset included a five-rod Mindways Model 3 CT calibration phantom within the field of view, enabling synchronous calibration using in-house software [17, 19]. The reference materials in the phantom span an equivalent density range from approximately –50 mg/cm³ to 375 mg/cm³ K₂HPO₄. Some scans contained partial phantom visibility, and full calibration certificates were not available.

### Finite Element Modeling

To estimate vertebral strength under compressive loading, we generated subject-specific finite element models from the calibrated CT scans and vertebral segmentations.

#### Mesh Generation

Vertebral body segmentations were converted into voxel-based hexahedral meshes using vktbone (1.1.0) and custom Python scripts based on VTK and SimpleITK. Images were resampled to 1.0 mm isotropic voxel size. Each voxel in the mask was treated as an 8-node hexahedral element (HEX8), and unconnected components were removed.

All models were solved using FAIM v9.0 (Numerics88) [41] with a convergence tolerance of 1e^-6^, a maximum of 30,000 linear iterations, and 10,000 plastic iterations. Convergence was monitored using the maximum relative displacement norm, a standard nonlinear FE analysis convergence criterion that monitors the relative nodal displacement between iterations [42]. All simulations successfully converged under these conditions.

#### Material Property Assignment

We generated two types of finite element models: one with linear elastic material behavior and one with an elastic-perfectly plastic constitutive law. For the linear model, the elastic modulus (E) was assigned voxel-wise based on the density–modulus relationship described by Kopperdahl, Morgan [22],

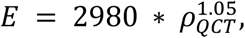

where ρ_QCT_ is the calibrated apparent density (g/cm³). For the nonlinear model, we defined a density-dependent yield stress

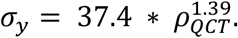

Cortical bone was not modeled explicitly, but implicitly accounted for through the naturally higher density values in cortical regions. Cortical bone segmentations included elements with apparent density above ∼1.0 g/cm³ and a 2 mm thick surface layer [24]. All bone material was assigned a Poisson’s ratio of 0.3. Simulations were conducted using 256 distinct density bins to approximate continuous material behavior.

#### Boundary Conditions and Load Application

To ensure consistent mechanical loading across subjects, we standardized vertebral alignment and applied uniform boundary conditions. Vertebral body masks were rigidly registered to a reference-aligned vertebra using an iterative closest point (ICP) transform. A single reference image (L4) was used and adjusted for smaller vertebra through principal component scaling. Superior and inferior surfaces were identified through an erosion-based morphological procedure. First, a 1-voxel-thick cortical shell was isolated based on density, and a single morphological erosion was applied using a 5×1×1 voxel kernel (z–y–x dimensions) to isolate flat superior and inferior surfaces suitable for load application and fixation.

Finite element simulations were conducted under uniaxial compression. The inferior surface of the vertebral body was fixed in all directions, while a uniform displacement was applied to the superior surface. Linear models were evaluated at 0.2% axial displacement, and nonlinear simulations were evaluated at 2% axial displacement. Boundary conditions were implemented using vtkbone (1.1.0). A 3-voxel-thick polymethylmethacrylate (PMMA) material was added to the superior and inferior surfaces, assigned a yield stress of 70.0 MPa and an elastic modulus of 2500 MPa and Poisson’s ratio of 0.3 to ensure consistent load distribution [43].

#### Force Estimation and Outcome Measures

We estimated failure load from linear models using two approaches. First, we evaluated three representative vertebrae with low, medium, and high density to determine the axial displacement at which the linear reaction force (R.Force) most closely matched the nonlinear failure load at 2% deformation. This analysis identified 0.2% displacement as an appropriate threshold. We then applied this displacement across the full cohort and extracted the reaction force from each linear simulation at this point. Second, we implemented a stiffness-based method based on the work by Crawford, Cann [18], where failure load (F.Load) was calculated as

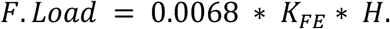

Here, K_FE_ represents the stiffness of the model (reaction force divided by applied displacement), and H is the vertebral height measured from the segmented mesh. This formulation approximates vertebral strength by assuming column-like mechanical behavior under axial loading.

For the nonlinear model, the ultimate failure load was defined as the force at 2% deformation. Stiffness was calculated as the slope of the initial linear region of the force–displacement curve. Bland–Altman analysis and Pearson correlation coefficients were used to assess agreement between methods.

#### Analysis of Vertebral Strength Variation

To evaluate intervertebral strength differences, we applied the linear finite element model across all vertebral levels and individuals. Two normalization strategies were used. First, within-subject comparisons were performed between adjacent vertebrae to calculate strength ratios. These ratios were then used to construct a graph model, where pairwise vertebral strength relationships were aggregated across subjects to estimate each vertebral level’s strength relative to L1. Only vertebrae that were anatomically adjacent and free of fractures or foreign materials were used to calculate stepwise ratios. Second, a population-level normalization was performed by computing the mean L1 strength across all individuals and comparing each vertebral level’s strength to this reference. While the first method captures individual variation and is less sensitive to scan coverage, the second provides a direct cohort-wide comparison. Results were stratified by sex and summarized in reference maps as percent differences from L1, with negative values indicating lower strength compared to L1.

## Statistical Analysis

All statistical analyses were performed in Python using NumPy [44], SciPy [45], and statsmodels [45]. Agreement between calibration methods and modeling approaches was assessed using Pearson correlation and Bland–Altman plots. Figures were generated using Matplotlib [46] and Seaborn [47].

## Availability of Supporting Data and Materials

All derived data used in this manuscript, including curated metadata (e.g., vBMD values, vertebral stiffness, and strength estimates), as well as code to generate figures and instructions for applying the data in other analyses, and sample data are available at <u>github.com/Bonelab/spineFE-benchmark</u> under the GNU General Public License v3.0 (GPL-3.0). The full dataset, including density-calibrated and aligned CT scans in NIfTI format and corresponding segmentation masks for vertebral bodies and processes (based on VerSe’19), will be released upon final publication. Until then, it is available upon request. The original VerSe dataset is publicly available from the VerSe challenge organizers at <u>osf.io/nqjyw</u> under the Creative Commons Attribution-ShareAlike 2.0 license (CC BY-SA 2.0). We provide modified versions of the data, including calibrated images and new segmentations, under the same license. Model weights for the nnU-Net (<u>github.com/MIC-DKFZ/nnUNet</u>) models are available on Zenodo for segmenting vertebral bodies and spinous processes [48], and tissues for phantomless calibration [49]. Both models are released under the Creative Commons Attribution 4.0 International Non-Commercial (CC BY-NC 4.0) license. Code for performing density calibration is available at <u>github.com/Bonelab/Ogo</u> under the GNU General Public License v3.0 (GPL-3.0), and the open-source finite element solver FAIM is available at <u>bonelab.github.io/n88</u>.

## List of Abbreviations

BCT: Biomechanical Computed Tomography
CT: Computed Tomography
DXA: Dual X-ray Absorptiometry
FE: Finite Element
HU: Hounsfield Unit
ICP: Iterative Closest Point
PMMA: Polymethylmethacrylate
QCT: Quantitative Computed Tomography
vBMD: Volumetric Bone Mineral Density

## Declarations

<u>Ethics Approval and Consent to Participate:</u> The original VerSe19 dataset used in this study was collected under ethical approval obtained from the Technical University of Munich (Proposal 27/19 S-SR), and released under the Creative Commons Attribution-ShareAlike 2.0 license (CC BY-SA 2.0). The present study did not involve the collection of new human or animal data.

## Consent for Publication

Not applicable.

## Authors’ Contributions

**MW**: Conceptualization, Methodology, Software, Formal analysis, Visualization, Writing – original draft. **BEM**: Data curation, Validation, Visualization, Writing – review & editing. **SKB**: Conceptualization, Supervision, Funding acquisition, Writing – review & editing.

All authors read and approved the final manuscript.

## Acknowledgements

We gratefully acknowledge the VerSe19 dataset, which provided the foundation for vertebral segmentations used in this study. The VerSe19 data were made available through the MICCAI 2019 Vertebrae Segmentation Challenge and are a valuable resource for advancing research in spine imaging and analysis.

